# Can elevated plasma Adiponectin and Ghrelin counteract inflammation in the aging heart?

**DOI:** 10.1101/2023.06.11.544501

**Authors:** Harsika Nahar, Shivanshu Chandan

## Abstract

The adaptability of the heart helps in sustaining its function under severe pressure overload conditions, including myocardial infarction and heart failure. Immune response and inflammatory changes are among the adaptive changes the heart relies on when challenged with stress or pressure stimuli. However, the immune system homeostasis declines with advancing age and increases the susceptibility to develop heart failure. Dissecting the inflammatory changes associated with age could develop novel rejuvenating therapies for an aging population. The older mice show tremendous cardiac adaptations with advancing age. However, how the old heart adapts and survives the chronic stress that increases with age are unclear. The potential involvement of inflammatory alterations in older heart has not been recognized previously.

We performed a screen of genes and proteins from RNA-seq and proteome profiles for regulators of cardiac inflammation in the old heart. We identified several pro-inflammatory and anti-inflammatory factors that belong to several immune response pathways. The inflammatory mediator plasma leptin levels increase at 3 months and decrease in the 18 months older mice. We found that the activated inflammatory gene program is associated with reduced left ventricular ejection fraction and vice-versa in the old mice. We also observed that elevated plasma levels of adiponectin and ghrelin are associated with reduced inflammatory molecules, including leptin, in these animals. We speculate that the induction of adiponectin and ghrelin secretion and downregulation of leptin secretion appears to encounter the elevated inflammatory gene program observed in the aging heart.

## Introduction

The heart beats approximately 3 billion times throughout 80 years lifespan; It adjusts the contractile function to sustain the altered oxygen demand. Several aetiologies including myocardial infarction, myocarditis, endocarditis, and arrhythmia hinder the heart’s ability to adapt under pressure overload conditions leading to heart failure.

Notably, activation of neurohormonal and sympathetic system dominate animal research and clinical research on heart failure. Altering these pathways using drugs including beta-blockers, ACE inhibitors, and angiotensin receptor blockers improved outcomes, especially in patients with left ventricular dysfunction or heart failure with reduced ejection fraction (HFrEF). More than 50% of heart failure admissions in hospitals are however heart failure with preserved ejection fraction (HFpEF). None of these drugs are effective in HFpEF patients. Interestingly, both HFpEF and HFrEF patients commonly have an association of adverse clinical outcomes and elevated levels of pro-inflammatory cytokines. The magnitude of pro-inflammatory changes is lesser in patients with heart failure than what would be seen in acute infections or autoimmune disease suggesting the contribution of inflammation and immune response either in the maintenance or deterioration of patients with heart failure.

Adipokines especially adiponectin, ghrelin, and leptin are the emerging mediators and therapeutic targets for managing inflammation and immune responses in several disease conditions including obesity, diabetes mellitus, and cardiovascular diseases. Adiponectin, adipose tissue-derived plasma protein regulates glucose level, lipid metabolism, and insulin sensitivity. Elevated plasma adiponectin is associated with increased obesity-induced endothelial dysfunction and hypertension, and protects against atherosclerosis, myocardial infarction, and diabetic cardiomyopathy. Lack of adiponectin is associated with increased endoplasmic reticulum (ER) stress, increased oxidative stress, increased proinflammatory responses, and decreased protective cytokines in the heart. Plasma leptin and its counterpart ghrelin levels are associated with inflammatory cytokines in patients with inflammatory bowel disease and pulmonary tuberculosis (TB). Despite the beneficial effect of leptin in diabetes mellitus type 2 and obesity, its elevated serum level is linked to increased oxidative stress, inflammation, thrombosis, arterial stiffness, and atherogenesis. Ghrelin is an endogenous ligand for groyoungh hormone secretagogue receptor (GHS-R), which regulates food intake, energy expenditure, and groyoungh hormone secretion. Notably, immune system activation and chronic inflammation are accompanied by reduced body mass and appetite suggesting crosstalk between immune and neuroendocrine systems through shared ligands and signaling pathways. The progression of these paradigmatic adaptations in the older age is also less understood. We observed an association of plasma adiponectin, ghrelin and leptin secretion and inflammatory changes in the old heart. The inflammatory mediators, plasma leptin levels are increased at 3 months of age and decreased in the 18 months older mice while adiponectin and Ghrelin levels are increased with age in these mice. The body weight, plasma insulin, and glucose level are unaltered in these mice. In conclusion, we discovered an association of adiponectin, ghrelin, and leptin and inflammatory mediators in the older heart.

## Results

### A). Global reprogramming of immunological changes especially inflammatory cytokines, chemokines, and immune cells mediated signaling in the LV tissue of older mice

Inflammation is widely recognized to have a role in the development of variety of diseases associated with aging especially in cardiovascular disease. To define the immunological changes in the OLD heart associated with aging, we performed RNA sequencing (RNA-seq) using left ventricular (LV) tissue of 3 months (3M) and 18 months (18M) OLD mice. Using a 1.3-fold cutoff and false discovery rate (FDR) <0.05 threshold for inclusion, we identified approximately 2335 differentially expressed genes and ∼1727 genes in LV tissue of OLD mice at 3M and 18M of age respectively compared with YOUNG counterparts. Venn diagram revealed that there were ∼1588 uniquely expressed genes and ∼927 uniquely expressed genes identified in LV tissue of OLD mice at 3M and 18M respectively (Fig. 1A). Gene ontology (GO) analysis using uniquely expressed genes revealed an enrichment of immunological pathways in these mice.

**Figure 1.**
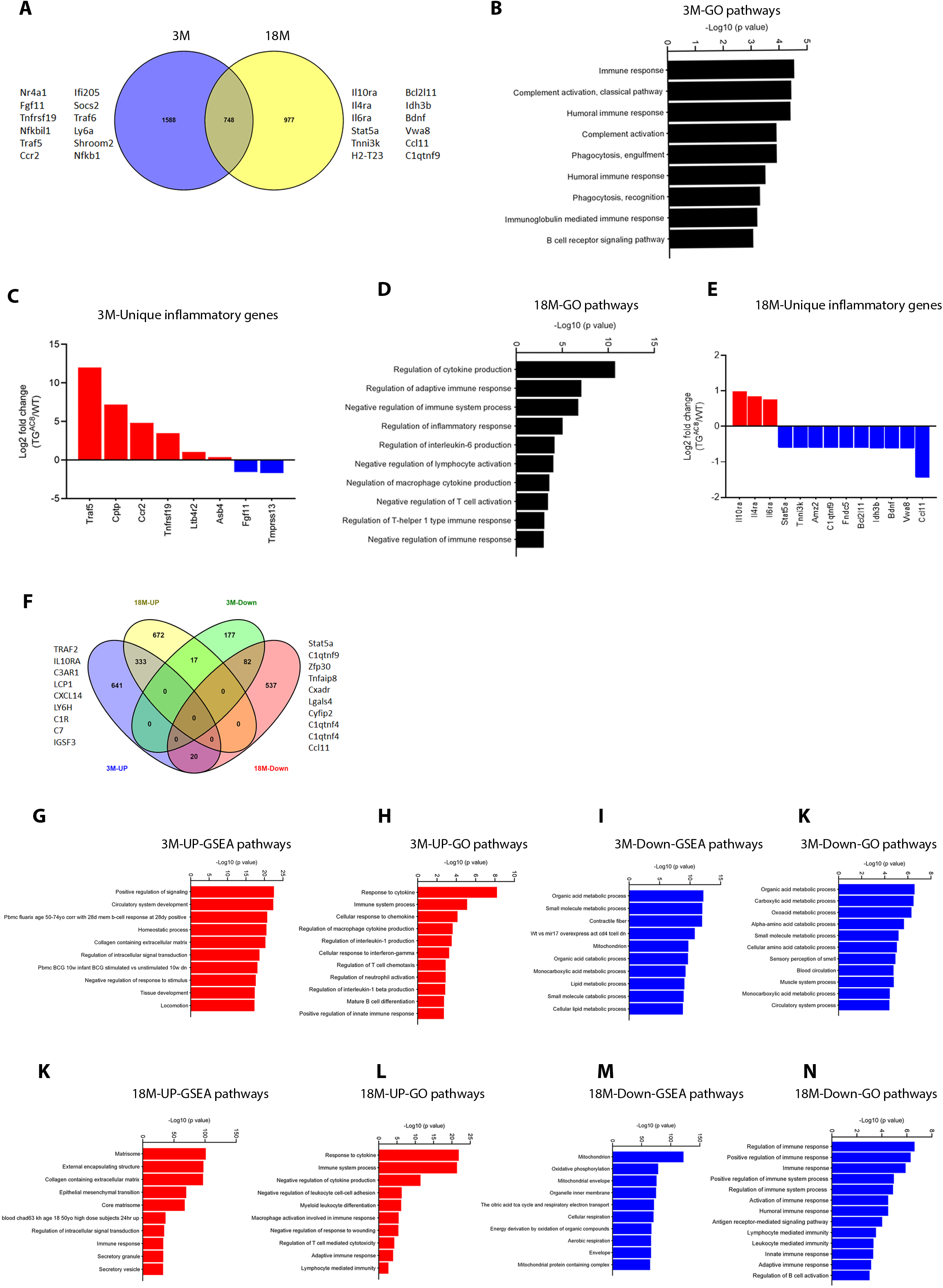

Among the enriched immune pathways, humoral immune response, complement activation and phagocytosis, and B-cell receptor signaling were observed at 3M; pathways negatively regulating the immune system process especially negative regulation of lymphocyte production, negative regulation of macrophage cytokine production and negative regulation of immune response dominated the immune pathways identified at 18M age group (Fig. 1B, 1C). Genes positively regulating the immune response especially Traf5, Cptp, CCR2, Tnfrsf19, Ltb4r2 were upregulated and Tmprss13 and Fgf11 were downregulated at 3M. Interestingly, genes which negatively regulate the cytokine signaling including IL10ra, IL4ra and IL6ra were upregulated and genes which positively regulate the immune system process including Stat5a, Tnni3k, Amz2, C1qtnf9, Fndc5, Idh3b, Bdnf, Vwa8 and Ccl11 were downregulated in 18M age group (Fig. 1D, 1E).

The Venn diagram analysis revealed that a distinct set of genes were upregulated and downregulated at both 3M and 18M age groups in the LV tissue of OLD mice (Fig. 1F). The Gene set enrichment analysis (GSEA) using upregulated genes at 3M revealed that BCG vaccine mediated immune response pathway was among the most enriched pathways at 3M in the OLD mice. Gene ontology (GO) analysis using upregulated genes also revealed that the cellular response to cytokines, IL-1 production, response to IFN-γ, T cell chemotaxis and B-cell activation pathways were among the top pathways upregulated at 3M in the OLD mice (Fig. 1G). The Gene set enrichment analysis (GSEA) using downregulated genes at 3M revealed that immune response pathways were not among the most enriched pathways at 3M in the OLD mice. Gene ontology (GO) analysis using downregulated genes also confirmed these findings (Fig. 1H).

The Gene set enrichment analysis (GSEA) using upregulated genes at 18M revealed that immune response pathway was among the most enriched pathways at 18M in the OLD mice. Gene ontology (GO) analysis using upregulated genes however revealed that the negative response to cytokines, leukocyte cell-cell adhesion, and negative response to wound healing pathways were among the top pathways enriched in the OLD mice (Fig.1I). The Gene set enrichment analysis (GSEA) using downregulated genes at 18M revealed that immune response pathways were not among the most enriched pathways in the OLD mice. Contrary to this, gene ontology (GO) analysis using downregulated genes at 18M revealed that the immune response pathways were among the enriched pathways in these mice (Fig. 1J). These results suggest a duel role of OLD on the immune response signaling in the OLD heart.

### B). Co-regulation of pro-inflammatory and anti-inflammatory factors in the OLD heart

Immune cell infiltration and aggressive cytokine response contribute to pathogenesis of the heart failure. In addition, immune cells especially cardiac resident macrophages facilitate electrical conduction of the heart by coupling to the cardiomyocytes in the atrioventricular node. We identified several pro-inflammatory and anti-inflammatory molecules in our RNA-seq and proteome data. Notably, the pro-inflammatory genes CD68 (macrophage marker), Stat3 (transcription factor regulating inflammation), Socs3 (suppressor of cytokine signaling) and the anti-inflammatory genes Mrc1, Mrc2, Tgm2 and Gpx3 were upregulated at 18M compared to the 3M in the LV tissue of OLD mice (Fig. 2A). Interestingly, proteome data also revealed that the pro-inflammatory proteins such as FABP4, Siglec-1, LYVE-1, Stat3, Stat6 expression was higher at 3M and anti-inflammatory proteins including Nid2, GGT1, Mrc1, Mrc2, Gpx3 and Igfbp5 expression was lower at 3M compared with 18M in the LV tissue of OLD mice (Fig. 2B). Western blot analysis confirmed the upregulation of RelA, a subunit of NFkB and CD68 protein expression at 3M in the LV tissue of OLD mice (Fig. 2C). The Quantibody inflammation array performed using plasma samples also confirmed that Lipopolysaccharide-induced CXC chemokine (LIX) and monocyte chemoattractant protein-1 (MCP-1) level were increased with age in both YOUNG and OLD mice The cytokines level was significantly higher at 3M and unaltered at 18M in the plasma of OLD mice compared to YOUNG counterparts (Fig. 2D). These observations suggest that the genes and proteins regulating pro-inflammatory signaling were increased at 3M; the anti-inflammatory mediators however dominated the inflammatory gene and protein signaling of 18M old OLD mice.

**Figure 2.**
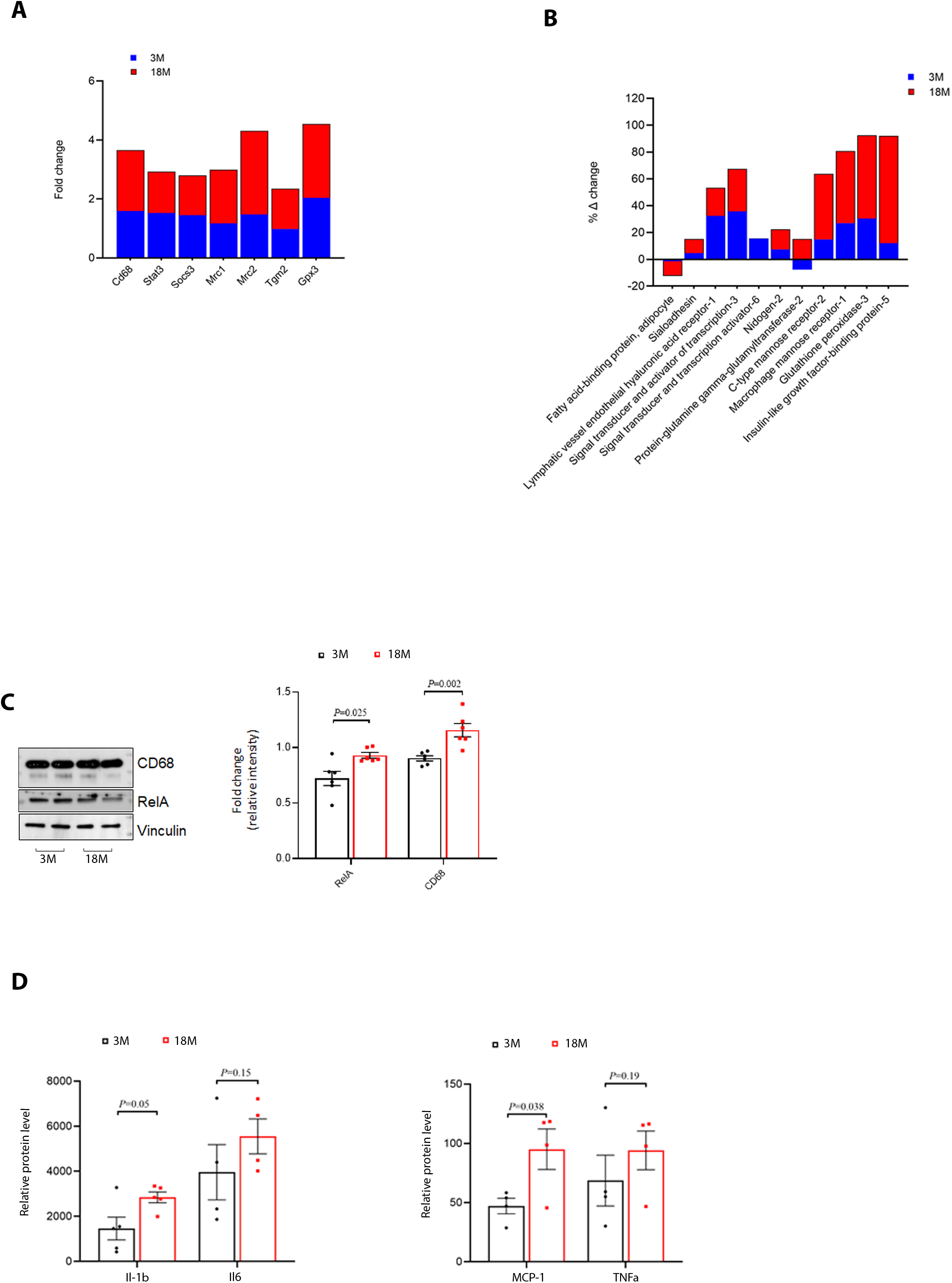

### C). The activated inflammatory gene program declines in the older mice with higher ejection fraction

Immune cells infiltrate the myocardium at gestation and contribute to its function throughout life. To define the inflammatory gene signature in the older mice and its association with ejection fraction (EF), we subdivided the 18M group into three different categories based on ejection fraction (1). OLD-Mid. EF Vs YOUNG-Mid. EF (2). OLD-Low EF Vs YOUNG-High EF (3). OLD-High EF Vs YOUNG-Low. The Venn diagram sketched using the RNA-seq data suggested a distinct gene expression pattern in all three different groups. There were several unique and common genes were observed in all the groups (Fig. 3A, 3B). Gene ontology (GO) analysis performed using upregulated genes in OLD-Mid. EF/YOUNG-Mid group revealed that immune system process, immune response and macrophage apoptotic process pathways were among the most enriched pathways identified. The GO analysis using downregulated genes of this group revealed the enrichment of pathways regulating cytokine production, macrophage CSF production, IL-β production, IFN-γ production and B cell negative selection (Fig. 3C, 3D). Gene ontology (GO) analysis performed using upregulated genes in OLD-Low. EF/YOUNG-High EF group revealed that negative regulation of immune response, B-cell chemotaxis, IL-8 production, TNF pathway, and negative regulation of adaptive immune response pathways were among the most enriched pathways identified. The GO analysis using downregulated genes of this group revealed the enrichment of pathways regulating positive regulation of macrophage differentiation, myeloid cell differentiation, neutrophil apoptotic process, and B-cell negative selection (Fig. 3E, 3F). Gene ontology (GO) analysis performed using upregulated genes in OLD-High EF/YOUNG-Low EF group revealed that negative regulation of B cell proliferation, B-cell proliferation and T cell differentiation pathways were among the most enriched pathways identified. The GO analysis using downregulated genes of this group revealed the enrichment of pathways regulating negative regulation of antigen-receptor signaling, neutrophil activation, leukocyte activation, cytokine production and B-cell negative selection (Fig. 3G, 3H). We then selected a set of inflammatory genes altered at 3M and compared their expression with all three 18M groups with different ejection fraction. Notably, a distinct set of inflammatory and immune response genes were observed in all three 18M groups with different ejection fraction compared to 3M (Fig. 3I-3N). In addition, we found that pro-inflammatory mediators such as Il1f9, Cx3cl1, and Lgals2 genes were upregulated in either OLD-Mid. EF Vs YOUNG-Mid. EF or OLD-Low EF Vs YOUNG-High EF and were not significantly altered in the OLD-High EF Vs YOUNG-Low group. Some pro-inflammatory genes such as Ilk, Il17rb and Il7 were downregulated in the OLD-Mid. EF Vs YOUNG-Mid. EF group and were not significantly altered in the OLD-Low EF Vs YOUNG-High EF and OLD-High EF Vs YOUNG-Low groups (Fig. 3J-3M). Interestingly, among these inflammatory mediators, anti-inflammatory markers genes Cd163, and CXCL16 were significantly upregulated in the OLD-High EF Vs YOUNG-Low group and were not altered in the Mid. EF Vs YOUNG-Mid. EF and OLD-Low EF Vs YOUNG-High EF groups. These results suggest that The inflammatory genes were distinctly expressed in in the 18M old OLD mice with different ejection fraction; the pro-inflammatory program genes were either not detected or were downregulated in the 18M old OLD mice with high ejection fraction.

**Figure 3.**
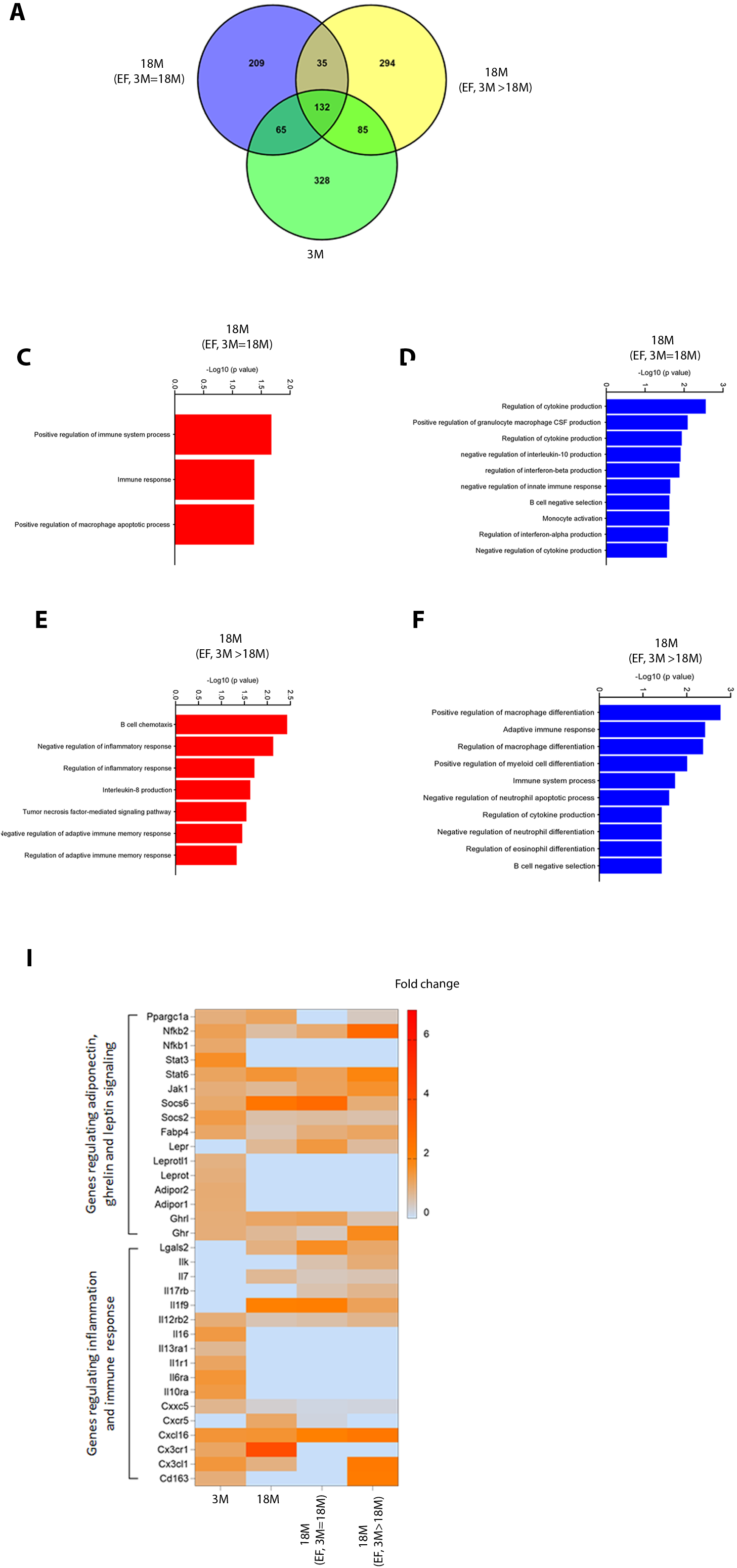

### D). The myocardial inflammation and innate immune response of OLD mice is associated with plasma levels of adiponectin, ghrelin and leptin

Adipokines alter chronic inflammatory state, cytokine level and thereby contribute to the pathogenesis of cardiovascular diseases. Consistent with these studies, we observed an up-regulation of genes (RNA-seq) regulating adipokine signaling including leptin, ghrelin and adiponectin signaling (Fig. 3I, Fig. 4A-4C). The set of upregulated genes including Socs2, Socs3, Fabp4, Jak1, Stat3, NFkB1, NFkB2 and Ppargc1b identified at 3M are involved in adipokine signaling (Fig. 4A); these genes were not significantly altered in the LV tissue of OLD-Mid. EF Vs YOUNG-Mid. EF and OLD-Low EF Vs YOUNG-High EF groups. Notably, inflammatory genes such as Jak1, Stat6 and NFkB2 were upregulated and leptin signaling genes including Lepr, Socs2, and Ppargc1a were downregulated in the LV tissue of OLD-High EF Vs YOUNG-Mid. EF group (Fig. 4A-4C). We next questioned if adipokine receptors such as leptin receptor, ghrelin receptor and adiponectin receptors are expressed in the LV tissue of these mice. Interestingly, Ghrl, AdipoR1, AdipoR2, Leprot1, and Leprotl1 were expressed at 3M and only Ghrl and Lepr were expressed in all the 18M groups with distinct EF (Fig. 4C). The leptin, adiponectin and ghrelin are secreted by adipocytes and released into blood. We measured the levels of these adipokines using plasma of both 3M and 18M mice. Leptin, adiponectin, ghrelin and levels were significantly altered with both age and genotype (Fig. 4D-4F). Plasma leptin level was significantly higher *(P<0*.*020)* in 3M and unchanged in 19M old OLD mice compared to YOUNG counterparts. Plasma leptin level was increased significantly *(P<0*.*008)* from 3M to 19M in the YOUNG mice and was unaltered in the OLD mice (Fig. 4D). Plasma adiponectin level was significantly higher *(P<0*.*010, P<0*.*002)* at both 3M and 19M old OLD mice compared to YOUNG mice. Plasma adiponectin level was increased significantly *(P<0*.*027, P<0*.*008)* from 3M to 19M in both YOUNG mice and OLD mice (Fig. 4E). Plasma ghrelin level was significantly higher *(P<0*.*001, P<0*.*001)* at both 3M and 19M old OLD mice compared to YOUNG mice. Plasma ghrelin level was decreased significantly *(P<0*.*006, P<0*.*001)* from 3M to 19M in both YOUNG mice and OLD mice (Fig. 4F). These results suggest an activation of adiponectin, ghrelin and leptin signaling in the OLD mice.

**Figure 4.**
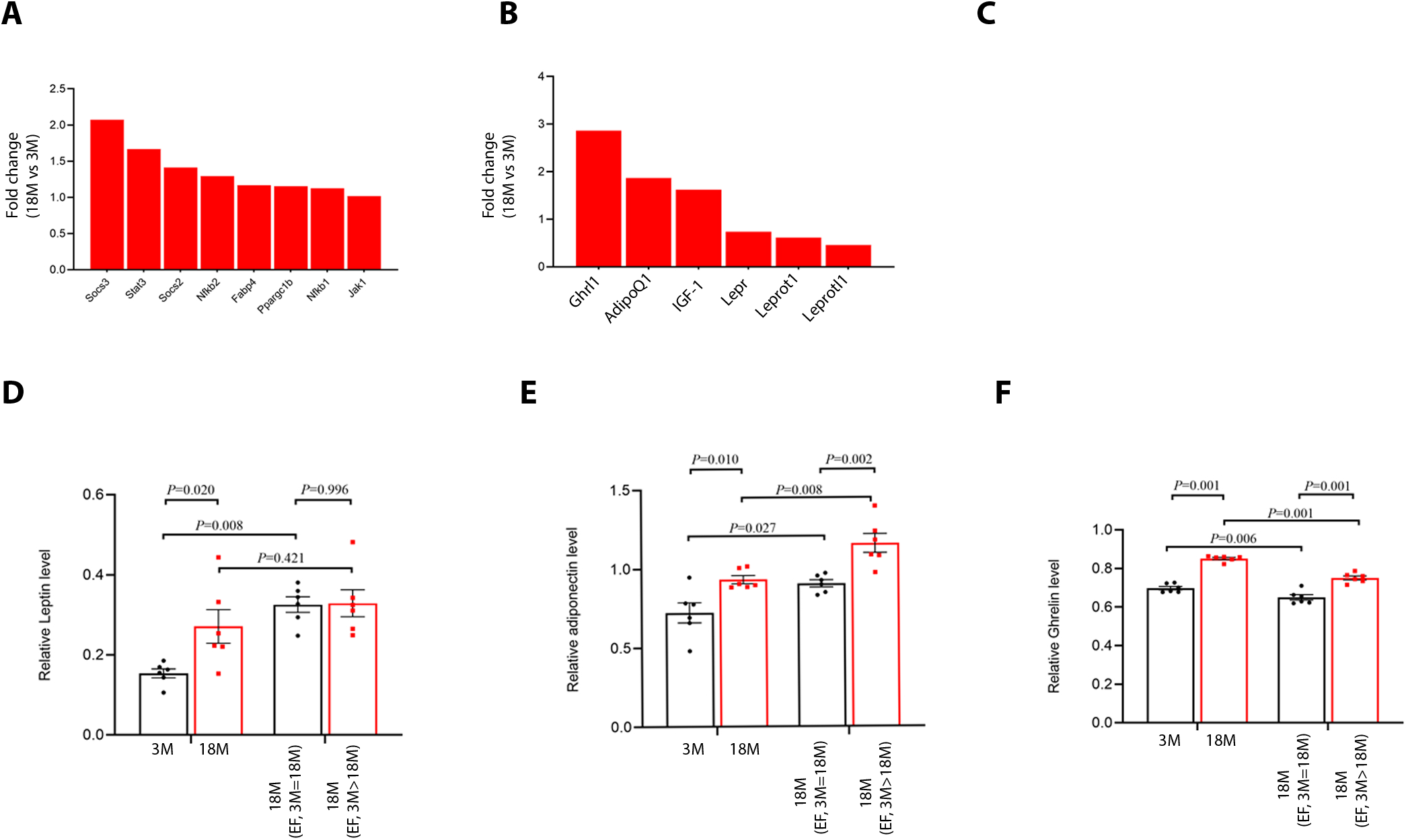

### E). The adiponectin, ghrelin, and leptin levels are not associated with weight gain and body fat in the OLD mice

Adipokines especially ghrelin and leptin regulate energy homeostasis, and in turn establish the nutritional status of an organism. The body weight was increased in both YOUNG and OLD mice with age; there was however no weight gain between YOUNG and OLD mice (Fig. 5A). The frailty index was increased with age in both YOUNG and OLD mice. There was no significant difference in frailty index in the OLD mice compared to YOUNG counterparts (Fig. 5B). The Micro–computed tomography (μCT) analysis revealed that the visceral fat was increased at 3M and unaltered at 19M in the OLD mice compared to YOUNG counterparts. There was no significant change in non-visceral fat at both 3M and 19M in the OLD mice compared to YOUNG mice. Notably, there was increase in visceral Fat, non-visceral Fat, Leg Fat and decrease in leg muscle with age in both YOUNG and OLD mice (Fig. 5B-5D). Metabolic activity measurement revealed that there was decrease in rate of oxygen consumption and carbon dioxide production with age in both YOUNG and OLD mice (Fig. 5E, 5F). The Respiratory Exchange Ratio (RER), body temperature, and food intake was not different in the OLD mice compared to YOUNG mice (Fig. 5G-5I). These results suggest that despite high level of leptin, and ghrelin at 3M in both YOUNG and OLD mice, there was no significant difference in weight gain, body fat and metabolic activity in these mice.

**Figure 5.**
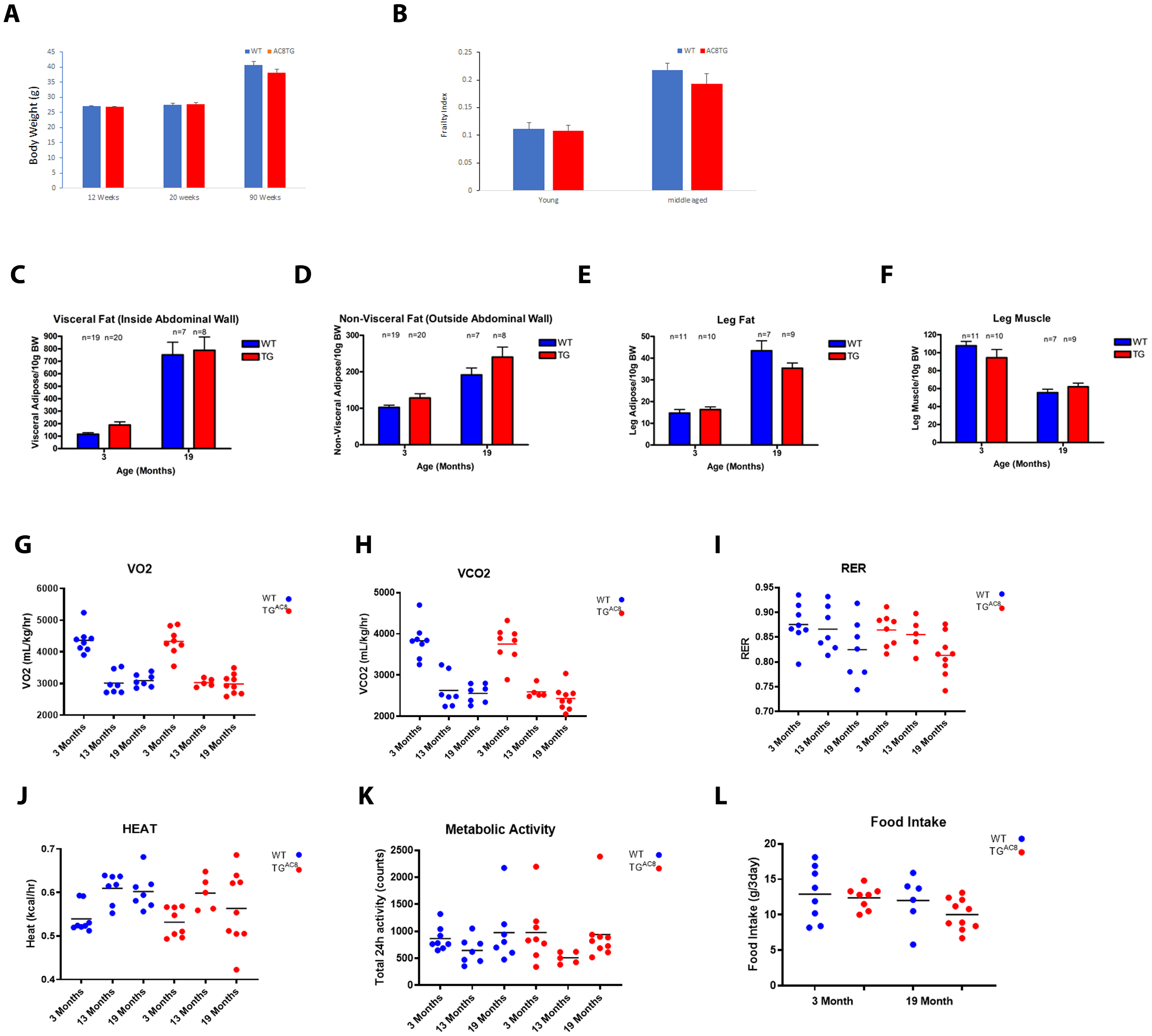

## Discussion

Activation of immune system and inflammatory gene program contribute to the pathological changes in aging heart. The potential involvement of adenylate cyclase 8 (OLD) in inflammatory gene program of the heart has not been recognized previously. Here, we present the data that implicates that elevated plasma adiponectin and ghrelin secretion can counteract inflammation in the aging OLD heart. We determined the Inflammatory gene signature of young (3M) and old heart (18M) with OLD gene. We performed a screen for regulators of cardiac inflammation in the OLD heart and identified pro-inflammatory and anti-inflammatory genes that belong to several immune response pathways. We found that the activated inflammatory gene program is associated with reduced left ventricular ejection fraction and vice-versa in the old mice. We also observed that elevated plasma levels of adiponectin, and ghrelin is associated with reduced inflammatory molecules including leptin in these animals. We specuate that cardiac specific OLD overexpression induce adiponectin and ghrelin secretion and downregulate leptin secretion to encounter the elevated inflammatory gene program observed in the aging heart.

### Declining pro-inflammatory cytokines, chemokines, and immune cells mediated signaling in the old OLD heart

With constant rise in aging population, it is important to understand the stress factors contributing to aging phenotypes. Inflammation especially immune cells infiltrate the heart and contribute to its contractile function; repair the scar left after myocardial ischemia, phagocytose apoptotic cells and produce soluble factors including cytokines and chemokines leading to acute inflammation (Aurora, A.B 2014; Esfahani, N.S 2021). A coordinated immune system is required for steady-state function of the heart; unfortunately, a constant decline in immune system function has been observed in elderly people which hinders the ability of the heart to adapt under pressure overload conditions including heart failure. We show that a unique set of anti-inflammatory factors and immune response pathways are upregulated in the aged OLD mice compared to YOUNG counterparts. A set of pro-inflammatory factors including Ccl11 and Stat5 are also downregulated in these mice as observed in previous studies (Finger, C.E., 2021). Aging is associated with infiltration of myeloid cells especially macrophages in the heart to suppress inflammaging mediated immuno-senescence (Hanna, A. 2020). Contrary to this, we observed an upregulation of proinflammatory factors in the young mice and anti-inflammatory factors in the old mice suggesting a reprogrammed inflammatory status of the OLD mice. The pro-inflammatory genes were upregulated in the old OLD mice with decreased cardiac function compared to YOUNG counterparts as observed in earlier studies (Triposkiadis, F., JACC; Esfahani, N.S., aging cell). We thus observed a distinct inflammatory gene pattern in the old OLD mice with different level of left ventricular ejection fraction. Previously, our lab demonstrated that despite constant activation of cAMP-PKA-Ca^2+^ signaling in SAN cells, OLD mice heart shows tremendous adaptations of limiting the additional activation of AC signaling; these adaptations were achieved by decreasing neuronal sympathetic input in the heart and ensured its survival (Moen, J.M et al 2019). We speculate that the increased cAMP-PKA-Ca^2+^ signaling contribute to inflammatory adaptations in these mice and help them in adaptations against severe Ca^2+^ overload as observed in earlier studies (Desvignes, L et al JCI). Also, overexpression of anti-inflammatory factors (Mrc1, Mrc2, Tgm2, Gpx3, GGT2, Nid2) in old age especially OLD mice with higher ejection fraction further establish the link between cardiac function and inflammation. Cardiac stem cell therapies underlie the beneficial effect of inflammatory response in rejuvenation of the heart after myocardial infarction (Vagnozzi, R.J Nature 2020; Bajpai, G., et al Circ. res 2019). Several lines of evidence indicate that adipokines especially adiponectin, ghrelin and leptin can regulate the pro- and anti-inflammatory signaling and energy homeostasis in several etiologies especially cardiovascular diseases (Van de Voorde, J Metabolism; Fuster, J.J., Circ res 2016). In line with these studies, we observed an increased systemic level of anti-inflammatory adiponectin and ghrelin molecules in young and old OLD mice especially old mice with increased cardiac function. Notably, pro-inflammatory peptide, leptin was significantly higher in young OLD mice and unaltered in old OLD mice. Interestingly, decreased leptin signaling genes especially Lepr, Socs2 in the old mice with higher ejection fraction strengthened the decreased pro-inflammatory signaling in these mice. We speculate that OLD mice at young age secrete leptin and other pro-inflammatory cytokine and chemokine factors to encounter the severe Ca^2+^ overload and cellular stress. Adiponectin and ghrelin are secreted to counterbalance the aggressive inflammation in the young and old OLD mice as observed in earlier studies (Cui, H., Nature Reviews Endocrinology 2017; Zheng, Y 2013).

OLD overexpression in mice led to aggravated cardiac dysfunction and heart failure with age and premature death compared to YOUNG counterparts (Mougenot, N 2019 CVR). This study however did not focus on the transcriptomic alterations especially inflammatory gene program in these animals.

